# Cryptic population structure and insecticide resistance in *Anopheles gambiae* from the southern Democratic Republic of Congo

**DOI:** 10.1101/2024.03.12.584605

**Authors:** Tristan P.W. Dennis, Poppy Pescod, Sonia Barasa, Louise Cerdeira, Eric R. Lucas, Chris S. Clarkson, Alistair Miles, Alex Asidi, Emile Z. Manzambi, Emery Metelo, Josue Zanga, Steve N. Nsalambi, Seth R. Irish, Martin James Donnelly, Fiacre Agossa, David Weetman, Francis Wat’senga Tezzo

**Affiliations:** Department of Vector Biology, Liverpool School of Tropical Medicine, Pembroke Place, Liverpool L3 5QA, UK; Pan-African Mosquito Control Association (PAMCA); Wellcome Sanger Institute, Hinxton, Cambridge CB10 1SA, United Kingdom; Unit of Entomology, Department of Parasitology, Institut National de Recherche Biomédicale (INRB/Kinshasa), 5345 Avenue De la Démocratie, Gombe, Kinshasa, Democratic Republic of the Congo; Faculty of Veterinary Medicine, National Pedagogical University, B.P 8815 Kinshasa, Democratic Republic of the Congo; Swiss Tropical and Public Health Institute (Swiss TPH), Kreuzstrasse 2, 4123 Allschwil, Switzerland; U.S. President’s Malaria Initiative (PMI) Evolve Project, Abt Associates, 6130 Executive Boulevard, Maryland, USA; Faculty of Medicine, School of Public Health, Department of Environmental Health, BP 834 KIN XI, University of Kinshasa, Kinshasa, Democratic Republic of Congo

**Keywords:** Insecticide resistance, WGS, malaria, Population genomics, *Anopheles gambiae* complex

## Abstract

The Democratic Republic of Congo (DRC) suffers from one of the highest malaria burdens worldwide, but information on its *Anopheles* vector populations is relatively limited. Preventative malaria control in DRC is reliant on pyrethroid-treated nets, raising concerns over the potential impacts of insecticide resistance. We sampled *Anopheles gambiae* from three geographically distinct populations (Kimpese, Kapolowe and Mikalayi) in southern DRC, collecting from three sub-sites per population and characterising mosquito collections from each for resistance to pyrethroids using WHO tube bioassays. Resistance to each of three different pyrethroids was generally high in *An. gambiae* with <92% mortality in all tests, but varied between collections, with mosquitoes from Kimpese being the most resistant.

Whole genome sequencing of 165 *An. gambiae* revealed evidence for genetic differentiation between Kimpese and Kapolowe / Mikalayi, but not between the latter two sample sites despite separation of approximately 800km. Surprisingly, there was evidence of population structure at a small spatial scale between collection subsites in Kimpese, despite separation of just tens of kilometres. Intra-population (H12) and inter-population (*F_ST_*) genome scans identified multiple peaks corresponding to genes associated with insecticide resistance such as the voltage gated sodium channel (*Vgsc)* target site on chromosome 2L, a *Cyp6* cytochrome P450 cluster on chromosome arm 2R, and the *Cyp9k1* P450 gene on chromosome X. In addition, in the Kimpese subsites, the P450 redox partner gene *Cpr* showed evidence for contemporary selection (H12) and population differentiation (*F_ST_*) meriting further exploration as a potential resistance associated marker.

## BACKGROUND

Malaria is a major burden in the Democratic Republic of the Congo (DRC), with 12% of all global malaria cases occurring in the country (World Health Organization, 2023b). Several *Anopheles* species are vectors of malaria in DRC, including the primary vector *Anopheles gambiae* and secondary vectors *An. funestus sensu stricto* (*s.s.*)*, An. coluzzii, An. arabiensis* and several uncommon or suspected vector species (e.g. *An. mouchetti, An. paludis, An. demeillioni*, etc.) (Irish *et al*., 2020). Resistance to pyrethroid insecticides, the main insecticide used on ITNs, the first line of defence against malaria, has been documented in *An. gambiae* in DRC; high frequencies of resistance-conferring mutations have been observed in the voltage-gated sodium channel (Wat’senga *et al*., 2018; The PMI VectorLink Project, 2021, 2022), and a recent selective sweep in the Cyp6aa / Cyp6p cluster of Cytochrome P450 genes has been implicated in enhancing metabolic resistance (Njoroge *et al*., 2022).

Malaria control programmes in DRC have depended heavily on the distribution of insecticide-treated bednets (ITNs) primarily using pyrethroids, with several nationwide distribution cycles over the last two decades contributing to a reduction in malaria incidence (Karemere *et al*., 2021; U.S. President’s Malaria Initiative, 2024). The scale of pyrethroid resistance across the African continent is prompting widespread deployment of synergists such as PBO and chlorfenapyr, which can help combat metabolic resistance (Hien *et al*., 2021). However, new generation nets are considerably more expensive than standard nets, and a better understanding of the distribution of resistance mechanisms would aid a more targeted distribution of these second generation nets (Gleave *et al*., 2018; World Health Organization, 2023a).

Despite the high impact of malaria in DRC there is a paucity of studies on its malaria vectors. The first iteration of the *Anopheles gambiae* 1000 genomes project (Ag1000G) increased understanding of the genetic variation underlying insecticide resistance in sub-Saharan Africa, but with a notable gap in the map covering the logistically complicated DRC (*Anopheles gambiae* 1000 Genomes Consortium *et al*., 2017). Subsequent additions to the Ag1000g dataset have facilitated analysis of genomic regions under recent selection in insecticide-resistant mosquito populations across Africa, demonstrating that the strongest selective sweeps in the genome include insecticide resistance genes (*Anopheles gambiae* 1000 Genomes Consortium *et al*., 2017; *Anopheles gambiae* 1000 Genomes Consortium, 2020). Here we describe the genomes of *An. gambiae* from collections made in three geographically distinct locations in southern DRC, phenotypically characterised for resistance to pyrethroids. We investigate the level of genomic differentiation at both fine and more distant scales, and identify both known and novel signals of selection likely linked to insecticide resistance in the region.

## RESULTS

Adult mosquitoes were collected for sequencing, and larval stages collected and reared to adulthood for phenotypic assays, at the sites shown in **Figure 1**.

**Figure 1.**
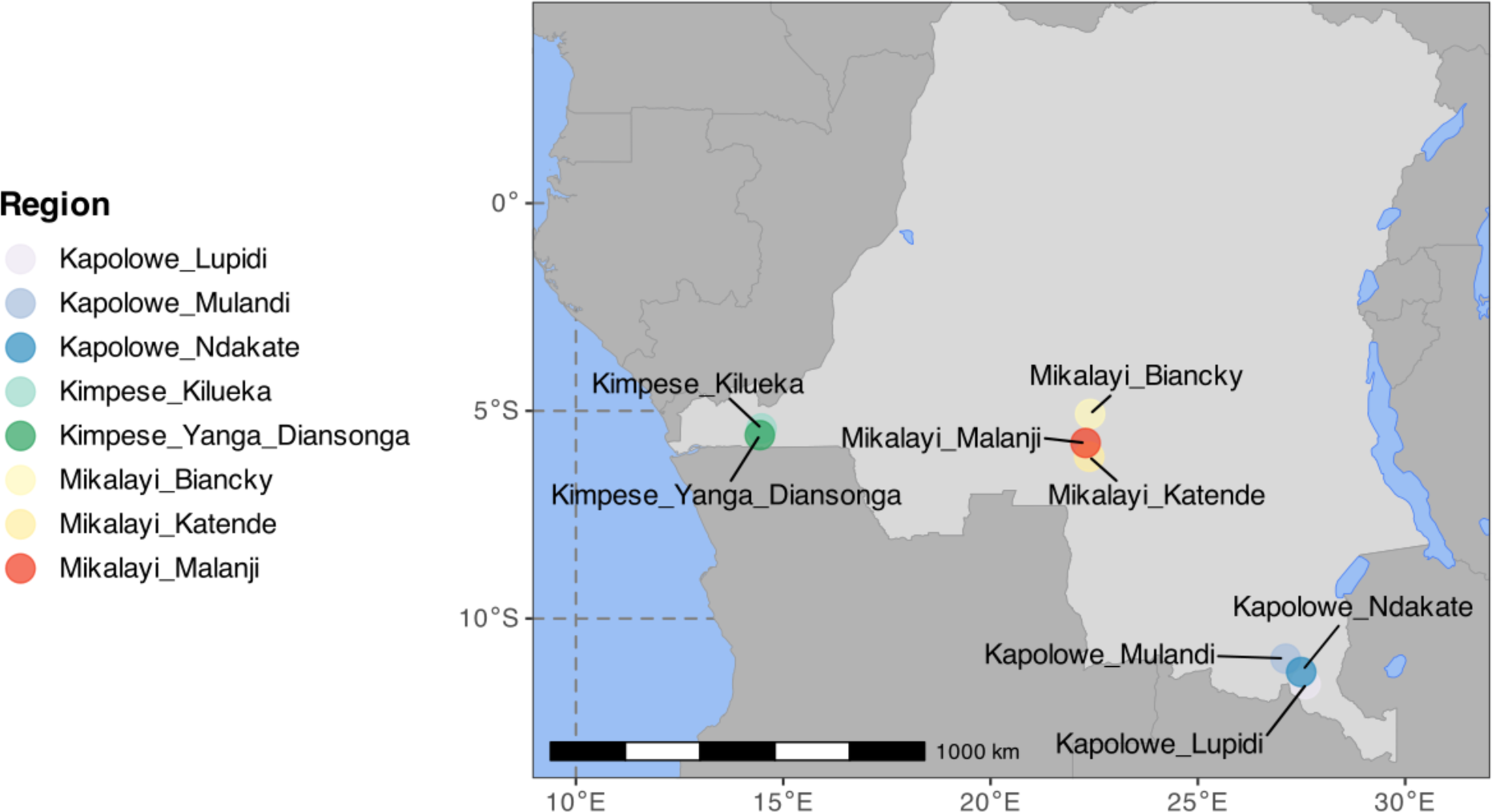
Map of the DRC showing the collection sites, where each circle represents a locality/village, coloured by the Province and location of each site

### Insecticide resistance bioassays

Larvae collected from each of the nine sites and reared to adulthood were tested against three insecticides at standard discriminating doses: alpha-cypermethrin (0.05%), deltamethrin (0.05%) and permethrin (0.75%). Bioassay results for *An. gambiae s.l.* are shown in **Figure 2** and given in full in Table S2. In all but one of the 27 site and insecticide combinations, mortality was less than 90%, indicating resistance according to WHO criteria (World Health Organization, 2022) - mortality after deltamethrin exposure was 91% in Bianki, Mikalayi, indicating suspected resistance.

**Figure 2.**
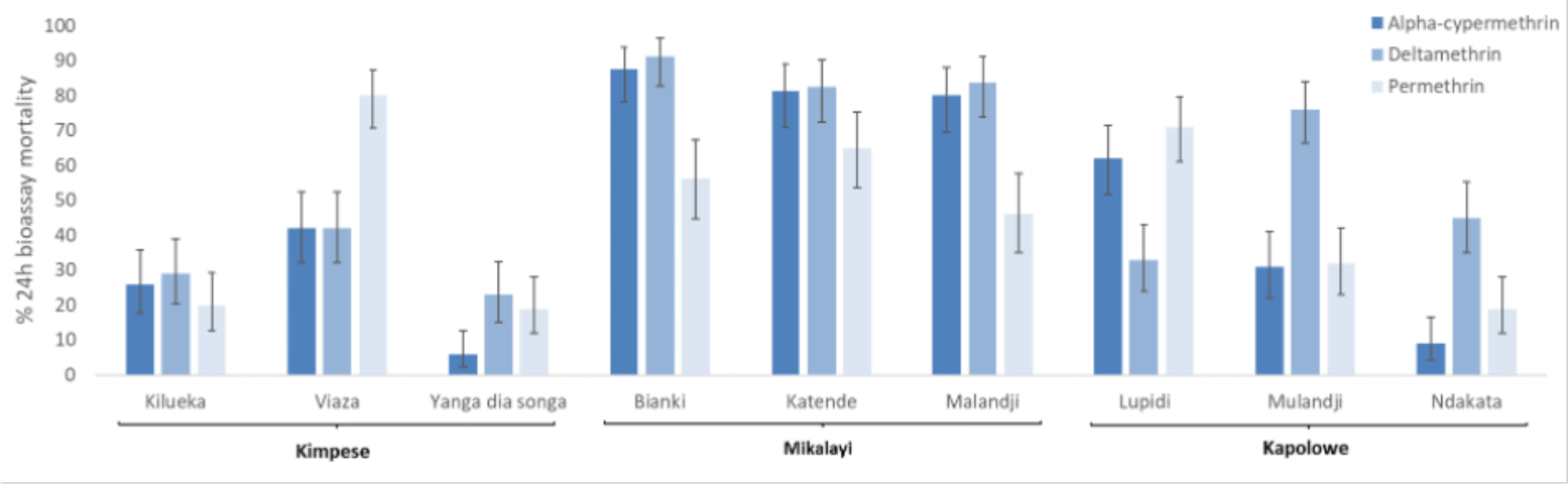
Pyrethroid insecticide resistance profiles for *Anopheles gambiae s.I* from the nine collection sites (three sites, each containing three sub-sites, 80-100 mosquitoes tested per site) assessed using WHO diagnostic bioassays. Error bars are 95% confidence limits.

Although all sites showed resistance to all three pyrethroids according to WHO criteria, there were significant differences between sites (Table S3); with Mikalayi showing least resistance and Kimpese showing strongest resistance (Table S4). Within sites there were also statistically significant differences in mortality to all three insecticides between some sub-sites; these differences were strongest between sub-sites in Kimpese, with Yanga Diansonga showing the highest resistance (Wald χ2 = 7.42, P = 0.006) and Viaza the lowest (Wald χ2 = 53.54, P < 0.001). The insecticide used also had a significant impact on mortality (Wald χ2 = 24.35, P < 0.001), with deltamethrin overall producing lower mortality than alpha-cypermethrin and permethrin (P < 0.001).

### Adult mosquito collection

Over the collection period, a total of 2,153 adult *Anopheles* mosquitoes were collected in Mikalayi, Kimpese and Kapolowe by pyrethrum spray catch with *An. funestus sensu lato* (*s.l*.) being the predominant species (64%), followed by *An. gambiae s.l.* (36%) and *An. paludis* (0.2%). The mean indoor resting density of *An. gambiae s.l.* was 4.8 and of *An. funestus s.l.* was 7.7 per house across the three provinces, with *An. gambiae s*.*l* mean densities of 6.1, 4.7 and 3.7 observed in Kapolowe, Kimpese and Mikalayi, respectively. The mean indoor resting density of *An. funestus s.l*. was 6.6, 14.2 and 2.3 in Kapolowe, Kimpese and Mikalayi, respectively. In all sites, the majority of *An. gambiae s.l.* (84.3%, 730/866) and *An. funestus s.l.* (82.5%, 1140/1382) collected indoors were blood-fed. One of the Kimpese sub-sites, Viaza, did not yield enough adult *An. gambiae* to be sequenced as part of the *Anopheles gambiae* 1000 genome project (*Anopheles gambiae* 1000 Genomes Consortium *et al*., 2017). Of the 237 samples sequenced, 165 passed quality control (QC) filtering. The samples were processed as part of the *Anopheles gambiae* genomics surveillance MalariaGEN Vector Observatory (VObs) project under sample set ID 1264-VO-CD-WATSENGA-VMF00164.

### Population structure and diversity

Principal component analysis (PCA) of all sequenced individuals (colour-labelled by collection site) for each chromosome is shown in **Figure S1**. At least three differentiation patterns are evident in the plots. On chromosomes X and arms 3L / 3R clear separation of the Kimpese (Kongo Central province) samples from those collected in Mikalayi (Kasai Central province) and Kapolowe (Haut Katanga province. Re-running the PCA as above, on chromosome 3L, on these two clusters (samples from Kimpese, and samples from Kapolowe/Mikalayi) (**Figure 3**), show no major hidden structure within them, though there is some separation between Kimpese Kilueka and Kimpese Yanga Diansonga (**Figure 3A**), and between Mikalayi and Kapolowe (**Figure 3B**). Separation within Kimpese is despite the sample sites being only approximately 20 km apart, the smallest pairwise distance among any of the sites and far less than the distance between Mikalayi and Kapolowe (approximately 800 km). As samples from sites within Mikalayi and Kapolowe districts appear to represent similar populations, these were pooled by district in subsequent analyses.

**Figure 3.**
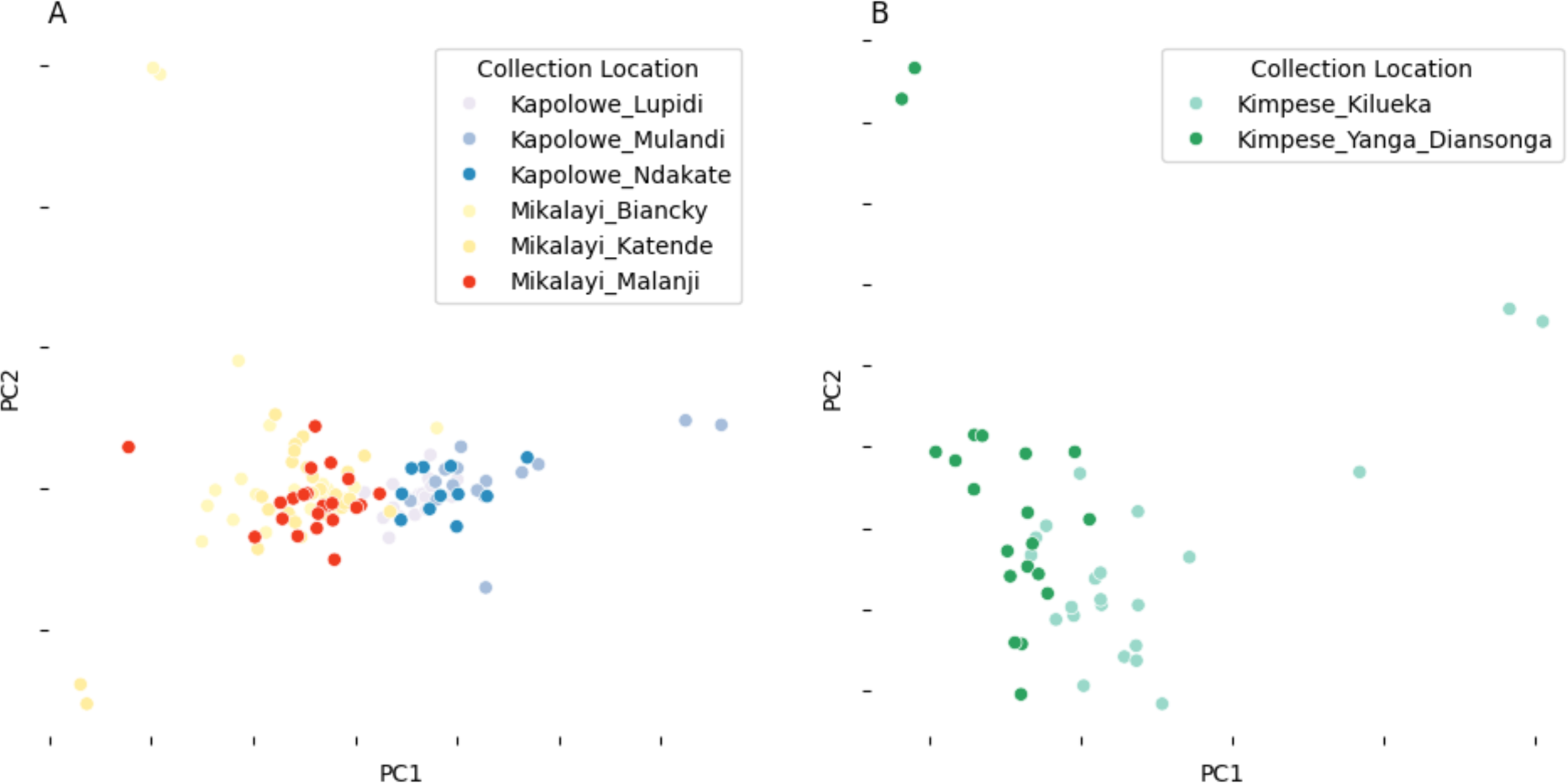
Principal component analysis (PCA) of samples from Kapolowe/Mikalayi (A) and Kimpese (B). PCA performed with genotypes from chromosome arm 3L. Points are coloured by collection location.

Further differentiation between sites is evident for chromosome 2, likely because of polymorphism in paracentric chromosomal inversions. This is most clearly evident for 2L, suggestive of 2La inversion karyotype variation. Separation is also evident on chromosome 2L PC2 and 2R PC2, also reflecting population structure among locations. In each case, more clearly for 2R, there is some evidence of separation of the Mikalayi and Kapolowe collection locations, likely suggesting differentiation in inversion frequencies. This contrasts with results for the other chromosomes (which lack polymorphic inversions in *An. gambiae*), indicating that differentiation between Mikalayi and Kapolowe is focal to inversions rather than genomewide.

### Genomic diversity

π, Watterson’s Θ (ΘW), and Tajima’s D were calculated and are shown in **Figure 4**. ΘW was highest in Kimpese-Kilueka and Kimpese-Yanga Diansonga, and lowest in Kapolowe (**Figure 4B**). π was highest in the two Kimpese cohorts compared to Mikalayi and Kapolowe. Tajima’s D was negative in all cases, and highest in Kapolowe. Kimpese-Kilueka had the lowest Tajima’s D. (**Figure 4C**)

**Figure 4.**
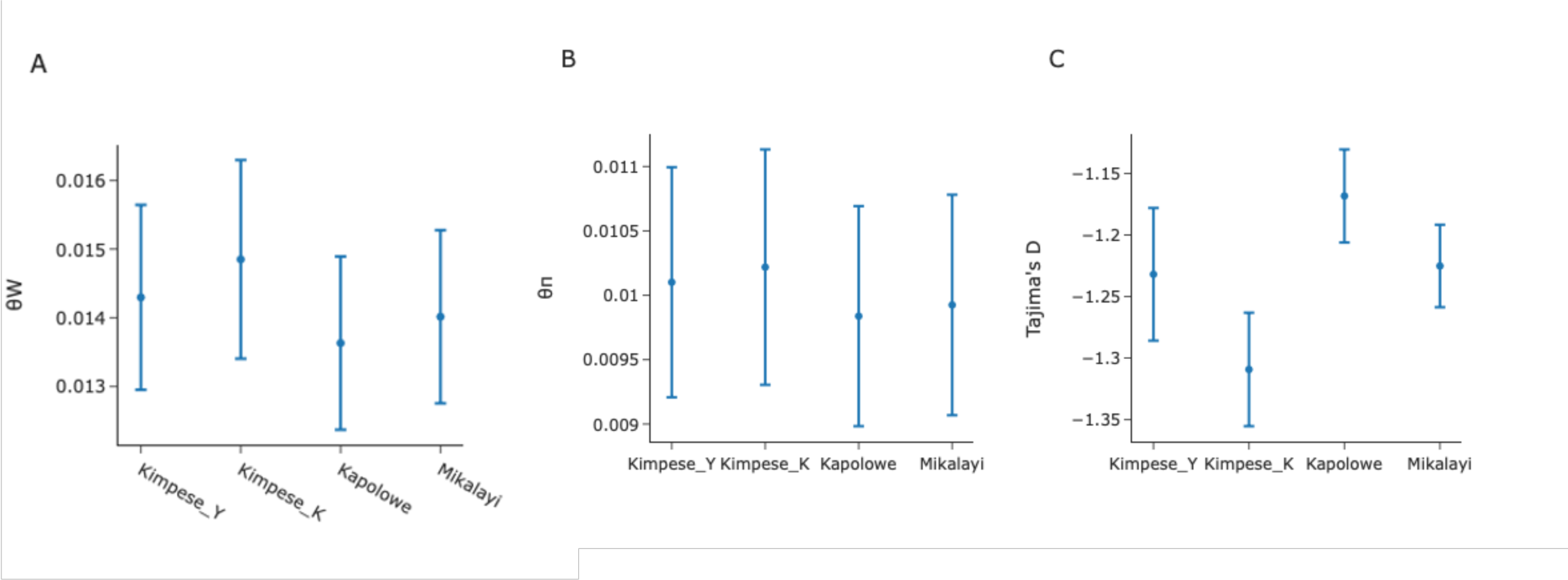
Genomic diversity statistics from (Kimpese_Y=Yanga Diansonga; Kimpese_K = Kilueka) with 95% confidence intervals.

### Genomewide scans for signatures of selection

The statistic H12 was used to investigate signals of selection within each population (**Figure 5)**. On chromosome 2, a pronounced peak around 28.5 Mb is present in Kimpese-Kilueka, Mikalayi and Kapolowe (**Figure 5**), and to a much lesser extent in Kimpese-Yanga Diansonga, in which the largest peak is located at around 40.5 Mb. The 28.5 Mb peak centres on a cluster of P450 genes, including the proven pyrethroid-metabolising genes *Cyp6aa1*, *Cyp6p3* and *Cyp6p4* (Ibrahim *et al*., 2018; Adolfi *et al*., 2019). The 40.5 Mb peak centres on a gene *AGAP003623* (long-chain acyl-CoA synthetase), which has no previous association with insecticide resistance. On chromosome 2L, the only clear area highlighted by H12 is a protracted region evident in all populations centred on the voltage-gated sodium channel (*Vgsc*), which contains well known resistance-conferring *kdr* mutations (**Figure 5**). The only obvious peak on chromosome 3RL is in Kimpese-Yanga Diansonga, and is a clear signal evident just before 40 Mb (**Figure 5**). This peak centres on *AGAP012290*, an unnamed gene with transmembrane neurotransmitter transporter activity, and six *Cyp9* subfamily P450 genes (CYP9L1-3, CYP9J3-5), which are not currently known resistance candidate genes. Chromosome X shows considerable heterogeneity in signals between populations. In Mikalayi (**Figure 5**), there are no obvious peaks, barring that proximate to the telomere (far left in **Figure 5**), which is present in all populations and contains an uncharacterised gene *AGAP000002.* In both Kimpese populations, a very high and broad peak centres on the well-established pyrethroid metabolising gene *Cyp9k1*, with a second strong peak evident near 8.8 Mb in Kimpese-Yanga Diansonga and to a slightly lesser extent in Kapolowe and Kimpese-Kilueka. The closest gene to this peak is *AGAP000500* (NADPH cytochrome P450 reductase, *Cpr*), which was identified in a previous genome wide scan as a strong potential candidate for resistance because of its essential role in all P450-mediated metabolism reactions (Lucas *et al*., 2023).

**Figure 5.**
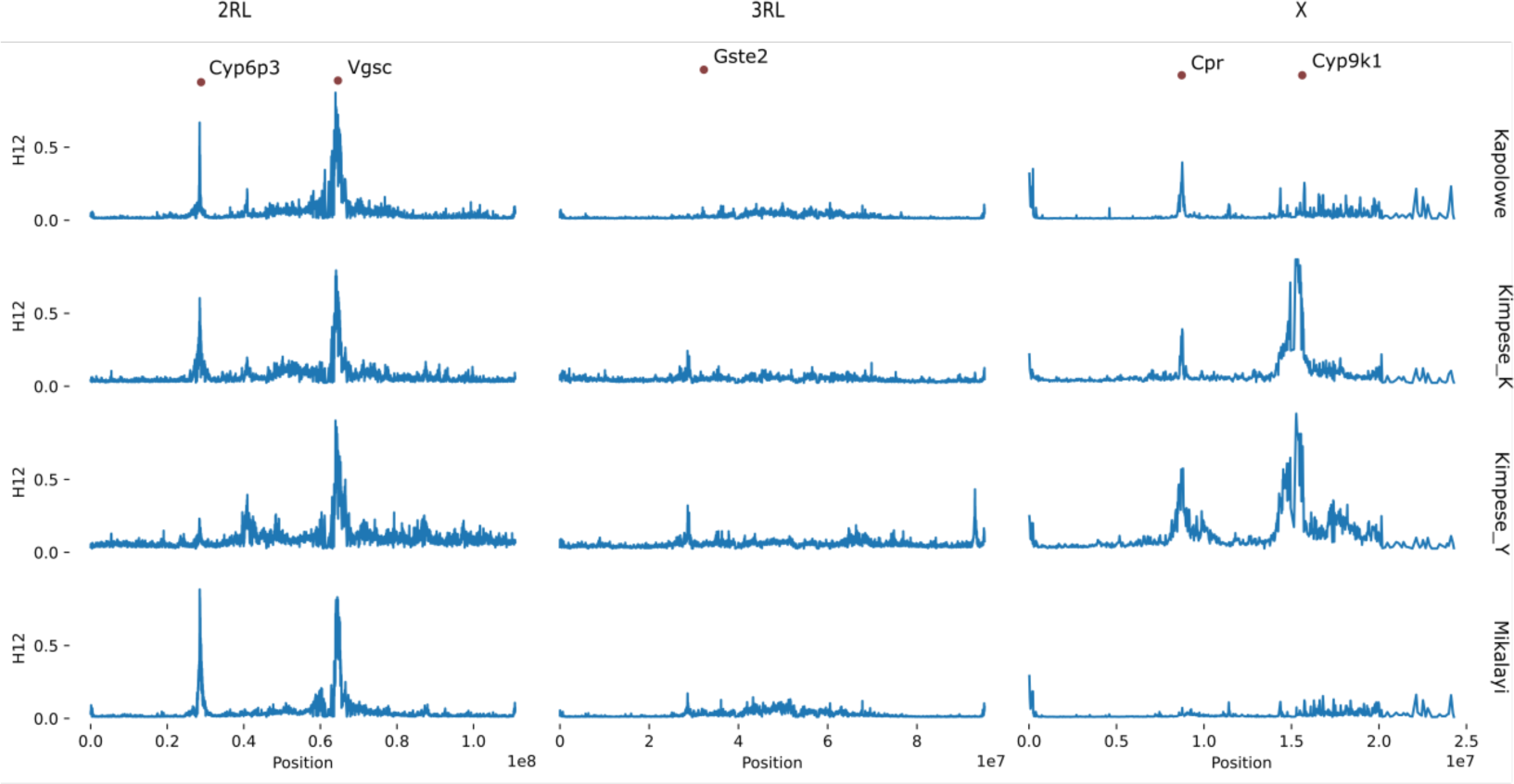
Genome-wide selection scan using the H12 statistic. X axis indicates position in a chromosome (plot columns), Y axis indicates H12 in a cohort (plot rows). Annotated points indicate IR genes of interest.

### Genomic comparison between populations using *F_ST_*

Genetic differentiation between the Mikalayi and Kapolowe populations, which showed differences in insecticide resistance profiles (see **Figure 2**) are shown in **Figure 6**. Moderate differentiation was seen in the *Vgsc* gene (2RL at approx 60Mb) between Mikalayi and Kapolowe, with stronger differentiation spanning the entire area of the 2La inversion visible, concordant with the differentiation between these sites on PC1 in **Figure 3**. Differentiation at the *Vgsc* between populations combined with a clear signal of selection (see **Figure 5**) may indicate population specific-haplotypes of *Vgsc,* or different frequencies of the same haplotype.

**Figure 6.**
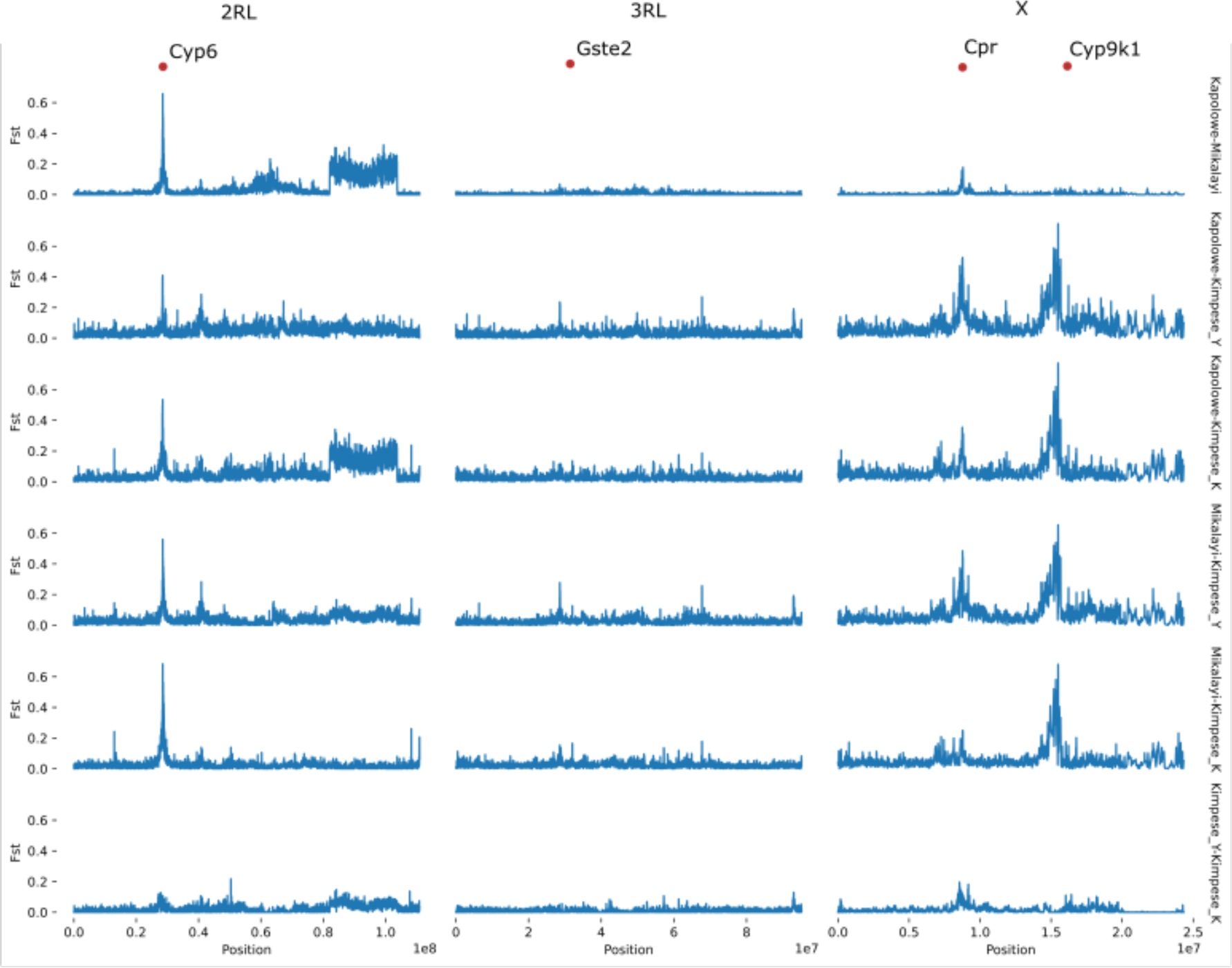
*F_ST_* comparison plots between population pairs. X axis indicates chromosomal position (chromosome denoted on the top of the plot), Y axis indicates *Fst*. Each row of plots denotes the labelled population pair. Genes of interest are highlighted on the top of the plot with red dots.

Differentiation on chromosome 2RL between Mikalayi and Kapolowe is dominated by an extremely strong, sharp peak centred on the *Cyp6* P450 cluster (**Figure 6**). Since H12 signals were similarly strong, this may indicate selection on different haplotypes in each population, or different frequencies of the same haplotype. A major peak area between 8.5 and 9 Mb dominates the *F_ST_* profile between Mikalayi and Kapolowe and the Kimpese sites for chromosome X, in proximity to the *Cpr* gene (**Figure 6**), and likely reflects a difference in the strength of H12 signal here between populations, which are present in Kapolowe but not in Mikalayi (**Figure 5**).

Differentiation between the two sites within Kimpese again highlights the *Cyp6* cluster on 2R (**Figure 6**), which was observed to show a much stronger H12 signal in Kilueka (**Figure 5**) than Yanga Diansonga (**Figure 5**). A second peak is also visible near 50 Mb. On chromosome X, differentiation around the *Cpr* gene is also evident, although far less pronounced than in comparisons with Mikalayi and Kapolowe (**Figure 6**).

Profiles of differentiation between the Kimpese populations on chromosomes 2L, 3R and 3L are less clear for the other chromosomes owing to relatively high baselines and lower peak *F_ST_*. Some differentiation of the 2La inversion region is present (**Figure 6**), but other peaks on each chromosome are generally not supported by multiple contiguous windows.

### Frequencies of known and candidate resistance markers

SNP data from were screened for non-synonymous (protein altering) mutations in the resistance-associated genes *Vgsc*, *Cyp4j5*, *Cyp6p4*, *Rdl*, and in the *Cpr* gene newly implicated via the signals of selection analyses, and can be seen in **Table 1** (more details in Table ST4). The G280S (previously G119S) resistance mutation in the acetylcholinesterase (*Ace-1*) target site gene was screened but was absent in the samples and is not shown.

**Table 1.** Frequencies of non-synonymous SNPs in insecticide resistance-implicated genes across the three study districts, with Kimpese district separated into its two sub-districts.

| Gene name | Substitution | Kimpese |  | Kapolowe | Mikalayi |
| --- | --- | --- | --- | --- | --- |
|  |  | Yanga Diansonga | Kilueka |  |  |
| *Vgsc* | R254K (G>A) | 0.85 | 0.77 | 0.92 | 0.77 |
|  | L995S (T>C) | 0.03 | 0.11 | 0.08 | 0.23 |
|  | L995F (A>T) | 0.97 | 0.89 | 0.92 | 0.77 |
|  | F1920S (T>C) | 0.13 | 0.09 | 0.00 | 0.00 |
| *Cyp4j5* | Q530E (G>C) | 0.73 | 0.43 | 0.82 | 0.40 |
|  | R509S (G>T) | 0.30 | 0.02 | 0.09 | 0.01 |
|  | L416V (G>C) | 0.13 | 0.00 | 0.00 | 0.00 |
|  | H415N (G>T) | 0.10 | 0.05 | 0.18 | 0.05 |
|  | I414N (A>T) | 0.10 | 0.05 | 0.18 | 0.05 |
|  | H377P (T>G) | 0.28 | 0.02 | 0.02 | 0.01 |
|  | N371D (T>C) | 0.10 | 0.05 | 0.04 | 0.12 |
|  | L354P (A>G) | 1.00 | 0.95 | 0.96 | 0.99 |
|  | I243T (A>G) | 1.00 | 1.00 | 0.99 | 1.00 |
|  | K216E (T>C) | 0.13 | 0.11 | 0.09 | 0.18 |
|  | A211T (C>T) | 0.90 | 0.70 | 0.93 | 0.76 |
|  | D177N (C>T) | 0.10 | 0.05 | 0.03 | 0.12 |
|  | W176R (A>G) | 0.08 | 0.11 | 0.21 | 0.05 |
|  | K175M (T>A) | 0.10 | 0.07 | 0.17 | 0.05 |
|  | V128I (C>T) | 0.45 | 0.23 | 0.71 | 0.19 |
|  | A69T (C>T) | 0.10 | 0.07 | 0.21 | 0.10 |
|  | R63G (T>C) | 1.00 | 1.00 | 1.00 | 1.00 |
|  |  | Yanga Diansonga | Kilueka |  |  |
|  | M58I (C>T) | 0.10 | 0.07 | 0.08 | 0.19 |
|  | M58V (T>C) | 0.13 | 0.02 | 0.03 | 0.05 |
|  | A47P (C>G) | 0.50 | 0.27 | 0.75 | 0.19 |
|  | L43F (G>A) | 0.50 | 0.27 | 0.74 | 0.19 |
|  | W30C (C>G) | 0.10 | 0.09 | 0.21 | 0.08 |
|  | T17S (T>A) | 0.00 | 0.00 | 0.11 | 0.01 |
|  | P12L (G>A) | 0.13 | 0.02 | 0.00 | 0.00 |
|  | S11A (A>C) | 0.13 | 0.02 | 0.00 | 0.00 |
| *Cyp6p3* | D418E (G>C) | 0.35 | 0.61 | 0.00 | 0.00 |
|  | L292V (A>C) | 1.00 | 1.00 | 1.00 | 1.00 |
|  | S287T (C>G) | 0.08 | 0.02 | 0.76 | 0.11 |
|  | I158L (T>G) | 0.98 | 1.00 | 1.00 | 1.00 |
|  | D155N (C>T) | 0.30 | 0.30 | 0.00 | 0.00 |
| *Cyp6p4* | R473H (C>T) | 0.10 | 0.16 | 0.00 | 0.00 |
|  | I333V (T>C) | 0.10 | 0.05 | 0.75 | 0.10 |
|  | V177L (C>G) | 0.15 | 0.25 | 0.22 | 0.17 |
|  | L161F (C>G) | 0.35 | 0.61 | 0.00 | 0.00 |
| *Gste2* | T154S (G>C) | 0.90 | 0.80 | 0.71 | 0.75 |
|  | K92R (T>C) | 0.15 | 0.14 | 0.28 | 0.16 |
|  | G26S (C>T) | 0.15 | 0.20 | 0.35 | 0.25 |
|  | N3K (G>{C,T}) | 0.98 | 0.95 | 0.86 | 0.88 |
| *Cpr* | K75N (G>T) | 0.00 | 0.00 | 0.51 | 0.01 |
|  | T319N (C>A) | 0.70 | 0.30 | 0.00 | 0.01 |
|  | A583S (G>T) | 0.18 | 0.41 | 0.00 | 0.01 |
| *Cyp9k1* | D350E (G>T) | 0.00 | 0.00 | 0.23 | 0.06 |

The most common mutation in *Vgsc* was 995F, which was present at high frequency in all populations, though notably lower in Mikalayi, which showed the highest frequency of the other well-known *Vgsc* mutant 995S; notably, the wild-type L995 was absent. Other resistance-associated non-synonymous SNPs in *Vgsc*, such as N1570Y, were absent; the only other present was R254K, which has previously been observed in close linkage with L995F but with an unknown effect on phenotype (Clarkson *et al*., 2021). This variation between populations in the balance between 995F and S likely explains the *Vgsc* differentiation (**Figure 6**) between Mikalayi and Kapolowe. Variants in the *Rdl* gene – a resistance-causing mutant (A296G) and linked, potentially compensatory variant (T345M) – were present at very low frequency, and the other known resistance-associated *Rdl* variant A296S was absent from the population (Grau-Bové *et al*., 2020).

*Cyp4J5* is used as a predictive resistance marker in East African populations using the L43F mutation as a diagnostic SNP (Weetman *et al*., 2018). Twenty four substitutions were detected in *Cyp4j5*; L43F is present in all populations analysed here, with frequencies ranging from 0.19-0.64 (**Table 1**). Four non-synonymous variants with unknown phenotypic effects were detected in *Cyp6p4* (**Table 1**), which is within a P450 cluster containing multiple duplications implicated in increased insecticide resistance (Lucas *et al*., 2023). Most notably, the L161F mutation was absent in both Kapolowe and Mikalayi while being present in Kimpese at 0.35 and 0.61 frequencies in Yanga Diansonga and Kilueka respectively.

The three non-synonymous polymorphisms (K75N, T319N and A583S) at the candidate gene *Cpr* identified from genome scans show variation between populations. Mikalayi, which notably is the location with the lowest pyrethroid resistance in bioassays (**Figure 2**), showed an extremely low frequency of each SNP (0.01 for each), whereas Kapolowe showed a relatively high frequency of the K75N mutant (0.51) but an absence of the others. In the two Kimpese populations, which showed higher overall resistance than Kapolowe and Mikayali (**Figure 2**), mutant T319N was at higher frequency in Kimpese Yanga Diansonga (which had the most resistant mosquitoes and the strongest H12 signal at *Cpr*) than Kimpese Kilueka (0.7 vs 0.3), whilst the A583S mutant showed the opposite pattern (0.18 vs 0.40) (**Table 1**). Noting also that differentiation in the F_ST_ scans was very high in this area of the genome between Kapolowe and Mikalayi and between the two Kimpese populations, these three mutants follow a pattern which could be compatible with conferring different levels of resistance.

Frequencies of known copy number variants (i.e. gene amplifications) are shown in Table 2 for resistance candidate genes (more details in Table ST5). Copy number variation (CNV) frequencies at *Gste2* are similar for the two Kimpese populations (0.65 and 0.59), and at lower frequency in Kapolowe than Mikalayi (0.26 vs. 0.61). Although these CNVs have been linked to signals of selection in other populations previously (Lucas *et al*., 2023), no convincing evidence was found from selection scans for peaks at the eGST cluster situated at approximately 28Mb on chromosome 3R in the populations surveyed here. Amplification of *Cpr* was only detected in Kapolowe, and at a similar frequency to the K75N mutant (0.57 and 0.51, respectively), suggesting possible association. Amplifications at the two adjacent *Cyp6aa* genes were absent or very rare in the Kimpese populations but at high frequency in Kapolowe and Mikalayi. Duplications of *Cyp6aa1* are associated with insecticide resistance, with one specific duplication (*Dup1*) co-occurring exclusively with the nearby *Cyp6p4* I236M mutation (Njoroge *et al*., 2022). While *Cyp6aa1* CNVs were present at frequencies of 0.65 and 0.93 in Kapolowe and Mikalayi respectively, the I236M mutation was absent in all sites (Table 2), suggesting a different duplication to the *Dup1* identified in east Africa previously.

**Table 2.**
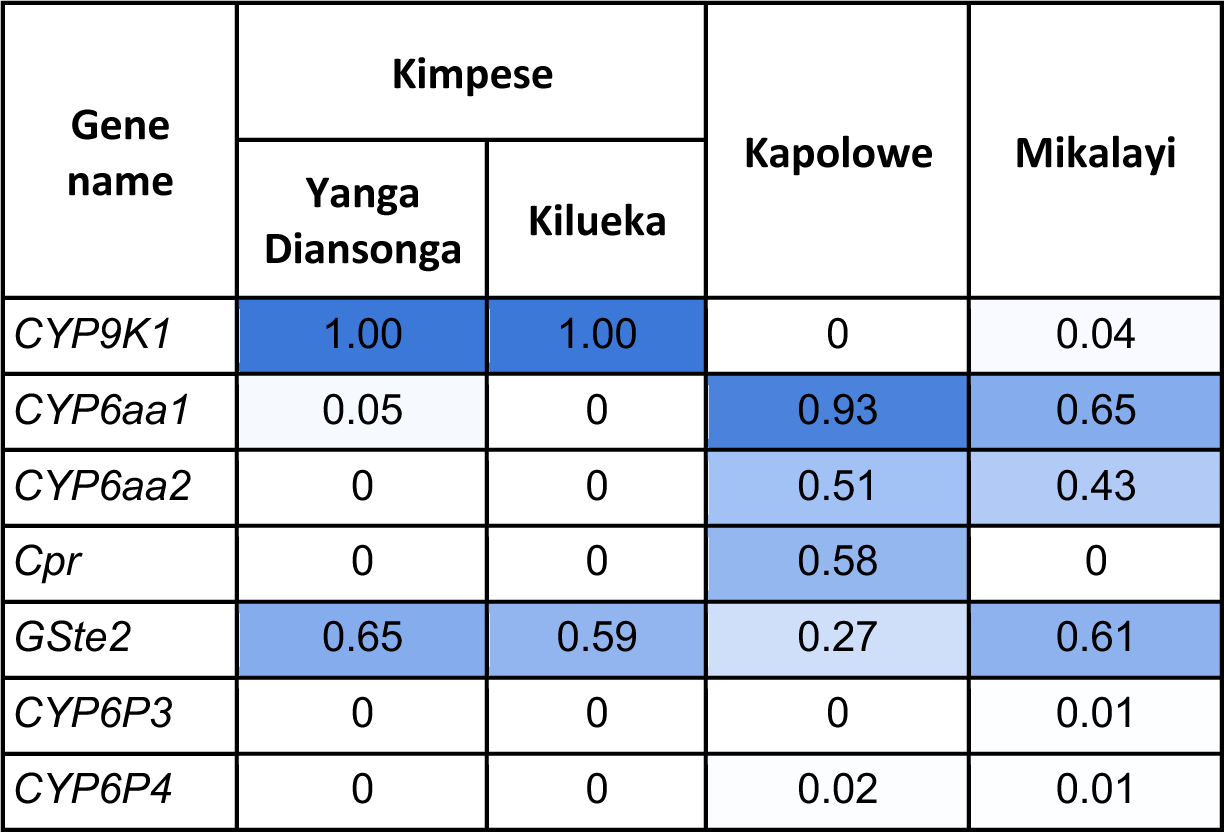
Frequencies of known copy number variants in candidate insecticide resistance genes.

*Cyp9K1* amplification was fixed in the Kimpese populations but absent from the other locations. This is of interest because a substantial H12 signal at *Cyp9K1* was seen only in the two Kimpese populations (**Figure 5**) and was absent from Kapolowe and Mikalayi (**Figure 5**), suggesting the duplication may be the target of selection in Kimpese where mosquito populations show highest pyrethroid resistance.

## Discussion

Despite the high burden of malaria in DRC, research on the major *Anopheles* vectors lags behind that from many other sub-Saharan African countries. The present study is the first population genomic investigations of insecticide resistance in *Anopheles gambiae* from the DRC. A striking finding was that of pronounced difference in population structure between the south-western sample site of Kimpese and the central and south-eastern locations of Mikalayi and Kapolowe, respectively. Despite comparable inter-sample geographic distance, *An. gambiae* from Mikalayi and Kapolowe grouped together in PCAs of chromosome 3 and X, and differentiation present on chromosome 2 was mostly attributable to inversion frequency variation. Marked differentiation between locations is uncommon in *An. gambiae*, with broad scale homogeneity across vast spatial scales (*Anopheles gambiae* 1000 Genomes Consortium *et al*., 2017; Schmidt *et al*., 2019; *Anopheles gambiae* 1000 Genomes Consortium, 2020). In this study, genomic differentiation appeared to be concentrated mostly in regions with genes associated with insecticide resistance and, in the case of *Cpr*, a gene with a mechanism for plausible resistance involvement. This difference is most striking between Kimpese: non-Kimpese sites, mirroring the trend of increased resistance in Kimpese compared to the other sites. Additionally, some differentiation within Kimpese is also concentrated in resistance-associated genomic locations, suggesting that barriers to gene flow may be raised by differential exposure to insecticides within Kimpese.

Resistance to pyrethroid insecticides was present in all sites, with strength of resistance varying depending on location, including between adjacent sub-sites. Several mechanisms which could underpin pyrethroid resistance were identified in the genome scans and candidate gene SNP analysis. Consistently strong H12 signals around *Vgsc* were detected in each location. High frequencies of the 995F mutation were detected with complementary presence of 995S mutants and complete absence of wild type leucine alleles, such that all individuals possessed a genotype expected to confer some level of knockdown resistance to pyrethroids (Reimer *et al*., 2008). Although, with the exception of the R254K mutation, which is present on the 995F haplotype background and has an unclear role in resistance (Clarkson *et al*., 2021), other variants were absent.

Additional proven and potential mechanisms of pyrethroid resistance were identified in the genome scans and candidate SNP analysis but were variable among populations. Each site showed H12 peaks centred on the *Cyp6* cluster of P450 genes on chromosome arm 2R although this was less pronounced in the Kimpese subsites Kilueka and Yanga Diansonga than in Kapolowe and especially Mikalayi. A strong, sharp *F_ST_* peak between Kapolowe and Mikalayi centred on this gene cluster suggests that the causal variation(s) underlying the H12 signal may have been different between populations. While *Cyp6p4* frequencies were very low in each population, moderate-to-high CNV frequencies affecting the *Cyp6aa* genes were identified, which are also associated with enhanced pyrethroid resistance (Njoroge *et al*., 2022; Lucas *et al*., 2023). *Cyp9k1* has previously been associated with pyrethroid resistance in genome scans (Main *et al*., 2015; *Anopheles gambiae* 1000 Genomes Consortium *et al*., 2017) and transcriptomic association studies, and has been shown to metabolise pyrethroids (Vontas *et al*., 2018). Here, the H12 signal at *Cyp9k1* was restricted to the two Kimpese sub-locations, which were also the most resistant to pyrethroids. Moreover, whilst non-synonymous variation was not found in *Cyp9k1*, a CNV was fixed in Kimpese, but absent or nearly so, in Mikalayi and Kapolowe. Whilst, as with many CNVs in *An. gambiae* detoxification-linked genes, conclusive evidence of causal relationships with resistance is lacking (Lucas, Rockett, *et al*., 2019), these results are consistent with a link between the *Cyp9k1* CNV and pyrethroid resistance.

A novel potential resistance mechanism was also highlighted in the H12 scans for chromosome X, with peaks of variable strength around the *Cpr* gene in three of the four locations. *Cpr*, also known as NADPH cytochrome P450 reductase (AGAP000500), is an essential electron donor in P450-mediated reactions, including insecticide metabolism. Increased *Cpr* expression has been implicated in insecticide resistance in *Culex* (Gong *et al*., 2022), and knockdown of the gene has been demonstrated to elevate permethrin susceptibility in *An. gambaie* (Lycett *et al*., 2006). Differentiation in *F_ST_* plots and frequency differences in non-synonymous variants were also evident among samples at this locus, with near absence of each of the three SNPs identified in Mikalayi, in which no H12 signal is present, intermediate frequency of a K75N mutant in Kapolowe, and also of the T319N and A583S mutants in the Kimpese subsamples. Correspondence of these variants with patterns of pyrethroid resistance across the populations and H12 signal suggest further exploration of their role as possible resistance markers is warranted. The final major H12 signal, which was present only in the Kimpese Yanga Diansonga subsite on chromosome 3L, covered several genes, including a cluster of *Cyp9j* P450s. *Aedes aegypti Cyp9j* P450s have been shown to metabolise pyrethroids (Stevenson *et al*., 2012) and *Cyp9j5* is also a weak metaboliser of pyriproxyfen (Yunta *et al*., 2016).

Beyond P450s and associated genes (*Cpr*), we also detected the presence of CNVs covering epsilon glutathione-S-transferases. Variants in, or overexpression of, these genes, particularly *GSTe2*, are most frequently associated with DDT resistance (Mitchell *et al*., 2014; Riveron *et al*., 2014; Adolfi *et al*., 2019) but have also been linked with resistance to pyrethroids (Riveron *et al*., 2014; Lucas, Miles, *et al*., 2019) and organophosphates (Adolfi *et al*., 2019). Though H12 signals that centre on the eGST cluster have been detected in multiple *An. gambiae* populations (*Anopheles gambiae* 1000 Genomes Consortium *et al*., 2017), we detected little evidence in our genome scans, suggesting that variation in these genes may be less important in southern DRC populations.

Overall our results highlight the value of newer generation nets, which either evade pyrethroid mechanisms by inclusion of an additional insecticide or bypass metabolic resistance mechanisms, such as *Cyp6* P450s, *Cyp9k1* and *Cpr* by inclusion of the synergist PBO. The results also highlight that geographic proximity may be a poor predictor of patterns of pyrethroid resistance and associated mechanisms, with distance an unreliable proxy for genomic differentiation, and that specific investigation of resistance in different localities will be required.

## Conclusion

Our study gives a first insight into the genomic landscape of resistance in samples from multiple highly-resistant populations from the DRC. The selective signals detected appear to focus on genes and variants known or plausibly linked to pyrethroid resistance; strong resistance to three types of pyrethroids was described in all study sites, with the caveat that the samples sequenced were not the same samples used for genomic analyses. The level of resistance and strong selection signals seen are consistent with the heavy use of pyrethroids via country-wide ITN distribution programmes in DRC. In very large countries with logistically-difficult travel, broad scale sentinel site-based surveillance of insecticide resistance is challenging and molecular surveillance of resistance, which can be applied to samples collected and preserved for later analysis, has particular value. Several of the regions highlighted in our analysis already have candidate DNA markers, or we have detected novel variants which can be tested for applicability as part of a diagnostic panel. Implementation of such panels, or, where possible whole genome sequencing based surveillance to monitor spatial variation to help guide ITN- type distribution decisions, and to assess changes over time in areas where particular products are distributed, represents a key aim for operational decision making and resistance management.

## METHODS

### Sample collection

Three study sites were chosen to cover a breadth of geographical locations across southern DRC: Kimpese, located in the Southwest, Mikalayi in the South-Central and Kapolowe in the South-East. Within each of these, three sub-sites (health areas) were identified for collections in 2019 and 2020. The locations and their geographical coordinates are provided in Table Supplementary 01. Adult *Anopheles* were collected in the early morning (06:00-09:00) from houses at each site using pyrethrum spray catches (PSC), according to the standard PMI- Vectorlink protocol (The PMI VectorLink Project, 2020). Female *Anopheles* were identified using keys (Coetzee, 2020) and stored for later analysis.

### Insecticide bioassays

Larvae were collected from various water sources at each site and reared locally to adults for insecticide resistance testing using standard WHO tube bioassay procedures (World Health Organization, 2016) with pyrethroid papers each at the diagnostic concentrations: deltamethrin 0.05%; alpha-cypermethrin 0.05%; and permethrin 0.75%. Between 80 and 100 3-5 day old adult females were tested for each subsite and insecticide in four tubes alongside a control tube containing 25 females. After a one-hour exposure period, females were transferred back to their holding tubes and maintained with sugar water for 24 hours when final mortality was assessed. 95% confidence intervals were calculated using the Newcombe method (Newcombe, 1998). Results were compared among locations and insecticides using a binomial generalised linear model.

### Whole Genome Sequencing (WGS)

A total of 253 collected adult *An. gambiae s.l* samples were sequenced on an Illumina HiSeq (150 bp paired-end reads) as part of the MalariaGEN Vector Observatory release 3.5, which comprises samples from eight of the nine collection sites; Viaza in Kimpese was dominated by *An. funestus*, providing too few *An. gambiae* for sequencing. Full details of library preparation, sequencing, alignment, SNP calling, CNV calling and phasing are detailed on the Ag1000G website (https://malariagen.github.io/vector-data/ag3/methods.html). Samples with coverage <10X on an Illumina HiSeq (150 bp paired-end reads) were removed. Sex was checked using the modal coverage ratio between chromosomes X and 3R (ratio between 0.4-0.6 = male, the ratio between 0.8-1.2 = female, other ratios would lead to sample exclusion).

### Read alignment and variant calling

Reads were aligned to the AgamP4 PEST reference genome using BWA (Li and Durbin, 2010), and indel realignment was performed using GATK version 3.7-0 (McKenna *et al*., 2010). Genotypes were called for each sample independently using GATK version 3.7-0 UnifiedGenotyper in genotyping mode, given all possible alleles at all genomic sites where the reference base was not ’N’. Additional details regarding thresholds and detailed information regarding read alignment, SNP calling, CNV calling and haplotype phasing are detailed on the Ag1000G website (https://malariagen.github.io/vector-data/ag3/methods.html). CartoDB library was used to plot the map layer, in Python and R.

### Population genomics

Population genomic analysis was performed using the malariagen_data *Ag3* API (MalariaGEN, 2023). Genetic diversity was assessed using S nucleotide diversity (π), which assesses the average number of pairwise nucleotide differences between DNA sequences in the sample; Watterson’s estimator (θ), which is a measure reflecting the number of segregating (i.e. polymorphic) sites in the sample; and Tajima’s D, which is calculated by comparing the two previous estimators, and for which negative or positive values indicate an excess or scarcity of rare alleles, respectively, relative to neutral model expectations. Genetic differentiation among individuals from each site was assessed using principal components analysis (PCA) run separately for each chromosome arm.

### SNP and CNV analysis

Frequencies of known mutants, either non-synonymous single nucleotide polymorphisms or copy number variants, were calculated for known or candidate resistance markers in a selection of genes, including the voltage-gated sodium channel (*Vgsc*) and gaba-gated chloride channel (*Rdl*) target sites, the P450 genes *Cyp4jJ5, Cyp6p4, Cyp6aa1, Cyp9k1*, and several Epsilon GSTs, as well as a relatively novel candidate, the P450 redox partner cytochrome P450 reductase, *Cpr*.

### Signals of selection analyses

Signatures of selection were assessed by genomewide scans using two statistical methods. The first, H12 (Garud and Petrov, 2016), was calculated using phased biallelic SNPs in 1000 SNP windows, using the garuds_h implementation in scikit-allel (Miles *et al*., 2023). H12 measures expected haplotype homozygosity when the two most common haplotypes within a single population are merged. Strong selection, primarily acting on one or two haplotypes, will increase the frequency and positivity of the H12 statistic. When present in multiple contiguous windows, this produces a pronounced peak in regions undergoing a selective sweep. The second statistic statistic, *F_ST_* (estimated using the method of Patterson in (Bhatia *et al*., 2013) via the moving_patterson_fst function in scikit-allel), compares allele frequencies in each window between populations; pronounced peaks suggest differentiation in that area of the genome, which may indicate differential selection between the populations.

### Data Summary

The authors confirm that all supporting data, code, and protocols are provided or referenced in the article or through supplementary data files. Code and data used in this study can be accessed at https://colab.research.google.com/drive/12dsseyUnndHqaMOzc_R2-lN7OW1nN3lI?usp=sharing. All sequence data generated in this study have been deposited in the European Nucleotide Archive (ENA) under project accession number PRJEB2141. Individual accession numbers are listed in **Table S1.**

## SUPPLEMENTARY FIGURES

**Figure S1.**
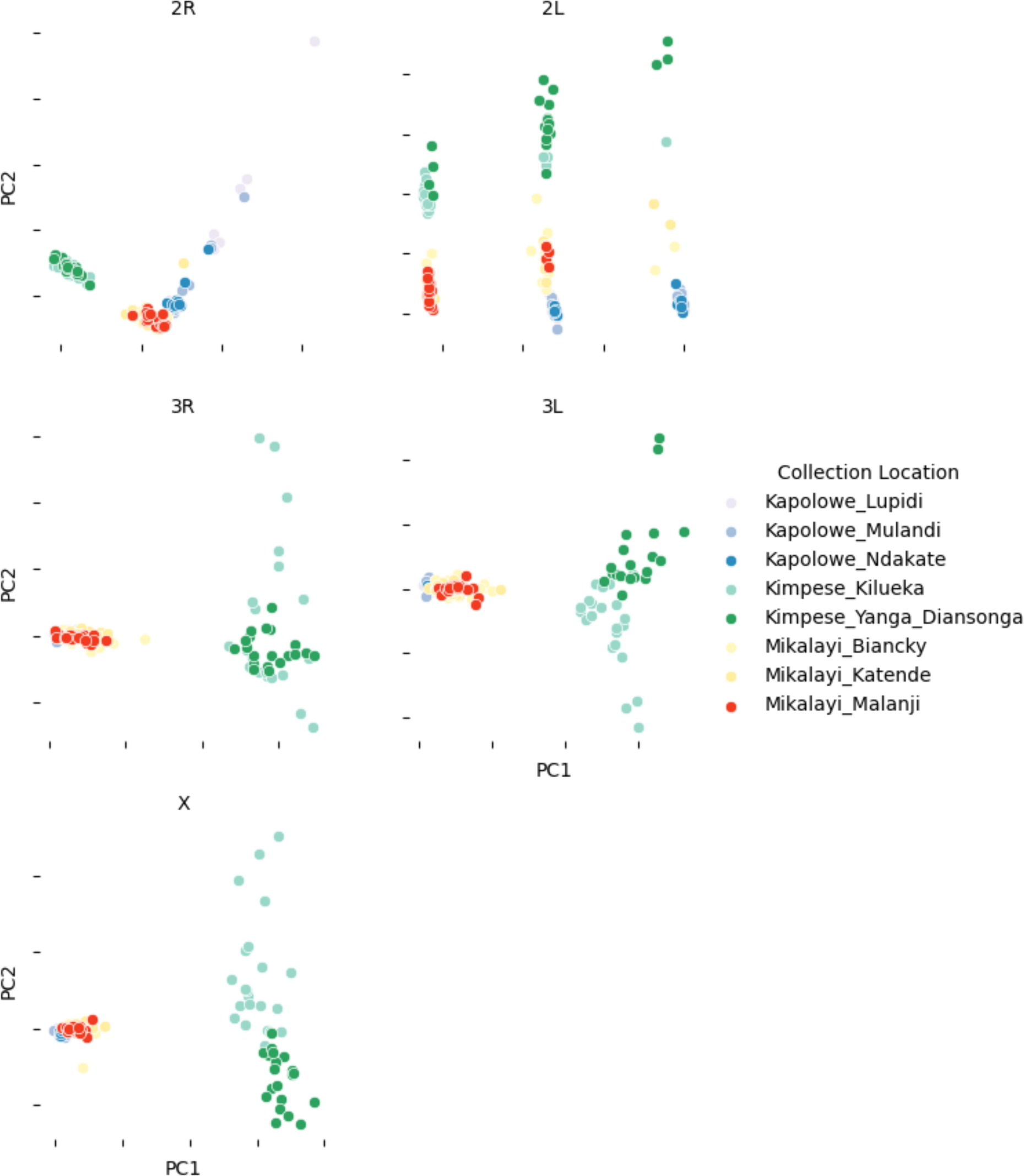
Principal component analysis (PCA) of all 165 sequenced *An. gambiae* samples. Plots are panelled by chromosome arm. Points, denoting individuals, are coloured by collection location, indicated in the key (which indicates collection district and collection site).

